# Polyphenol-rich fraction of Bergenia ligulata (Wall.) Engl. sensitizes colon and breast cancer metastasis in vivo: Evidence of cell-type specific mechanism

**DOI:** 10.1101/2024.10.06.616842

**Authors:** Samhita De, Suvranil Ghosh, Somesh Roy, Kuladip Jana, Subhas C Mandal, Mahadeb Pal

## Abstract

**Background:** Colon and breast cancer are one of the leading causes of mortality worldwide. Limited efficacies of present treatments highlighted an urgent need for improvement. Bergenia ligulata is known for its anticancer properties in Indian traditional and folk medicine though the molecular basis of its effects, particularly its anti-metastatic properties, is not well understood. Anti-prostate cancer activity of a LCMS defined polyphenol-rich fraction from *Bergenia ligulata* (PFBL) rhizome extract has already been published showing promising results in preclinical models.

**Purpose:** The present study aims to explore anti-cancer activity and anti-metastatic potentials of PFBL against colon and breast cancers in both in vitro and preclinical settings.

**Study design and methods:** PFBL alone or in synergy with standard chemotherapeutic drugs was tasted in CT26 and 4T1 subcutaneous solid tumors in BALB/c mice. The effect of PFBL was analyzed in terms of changes in tumor mass and different molecular marker levels. Anti-metastatic potential of PFBL was evaluated in CT26 and 4T1 lung metastasis model in mice focusing on the number of lung nodules and lung size.

**Results:** Our results evidenced that PFBL efficiently regressed both CT26 and 4T1-induced solid tumors alone and in combination with 5FU/doxorubicin without affecting the health of normal host. Notably, PFBL suppressed lung metastasis of 4T1 and CT26 cells in mice with great efficacy. Analysis of tumor and cell extracts suggested that colon cancer cells died by autophagy, while breast cancer cells majorly died by caspase-dependent apoptosis. PFBL actions involved AMPK-dependent inhibition of mTORC1 and subsequent increase in LC3II, PARP1, CDK4 and Cyclin D1 levels in both HCT116 and MCF7 cells; an elevation of intracellular ROS level was the major cause of death in both the cancer types. PFBL treatment also reversed the EMT marker expressions in in vivo and in vitro.

**Conclusion:** PFBL regressed colon and breast cancer metastasis through distinct mechanisms with little effect on healthy mice. The present study underscored PFBL as a novel anti-tumor and anti-metastasis candidate warranting further testing in clinical settings.

## 1. Introduction

Breast cancer and colorectal cancer (CRC) are two major causes of mortality and morbidity in the world. With more than 2 million new cases, breast cancer is the most prevalent cause of death in females, while CRC ranks as the 2nd major cause of cancer deaths with 1.9 million new cases claiming about 1 million lives in 2020 (Sung et al., 2021). Projections predicts that the global CRC burden to rise by 60% to 2.2 million new cases and 1.1 million deaths by 2030 (Sung et al., 2021). Additionally, cancer incidences were predicted to increase by 47% in the next two decade (Sung et al., 2021). It is forecasted that breast cancer burden will increase with 3.19 million new cases with 1.04 million deaths by 2040 (Arnold et al., 2022).

An increasing number of studies reporting significant anti-cancer activities in various plant extracts and compounds highlighted their potential use as chemotherapy agents alone or in combination with existing therapy regimens (De et al., 2023). *Bergenia ligulata*, locally known as “Pashanbhed”, is already known for its various medicinal significance, including anti-urolithic, anti-plasmodial, (Rajbhandari et al., 2003), antilithiatic (Aggarwal et al., 2014) anti-microbial (Agnihotri et al., 2015), anti-oxidant, and anti-inflammatory (Singh et al., 2022) properties. Recent studies have analysed the anti-cancer activity in *Bergenia ligulata* (Faheem et al., 2022; Ghosh et al., 2021). *Bergenia ligulata* silver nanoparticles showed anti-breast cancer activity in a p53 and ROS dependent manner (Faheem et al., 2022). We have demonstrated that PFBL, of defined chemical composition per LC-MS analysis, sensitized PC3 cells by robust upregulation of reactive oxygen species (ROS), and in PC3 cell-induced tumor xenograft in mice (Ghosh et al., 2021). Ghosh et al., (2021) demonstrated that this robust ROS production in the tumor involved of catalytic activation monoamine oxidase A (MAO-A) with simultaneous inactivation of NRF2 antioxidant response involving GSK3β. Notably, PFBL did not show any adverse effects on normal cells/healthy animals due to the elicitation of robust NRF2-dependent antioxidant response in them (Ghosh et al., 2021). These observations led us to analyse the efficacies of PFBL in metastatic model of breast and colorectal cancers in mice including the underlying mechanisms of action. This study uniquely indicated that PFBL inhibited these two types of cancer by distinct mechanisms.

## 2. Materials and methods

### 2.1. Cell culture

HCT116, MCF7, NKE, CT26 and 4T1 cells were purchased from ATCC and cultured as described by Ghosh et al., (Ghosh et al., 2021).

### 2.2. Polyphenol-rich fraction of *Bergenia ligulata* (PFBL)

LCMS characterised PFBL fraction extracted from *Bergenia ligulata* (Wall.) Engl. rhizome was used in this study as described (Ghosh et al., 2021).

### 2.3. Cell based analysis

Cell viability assay, Cell cycle analysis, and Apoptosis Assay was performed as described by Ghosh et al., 2021 (Ghosh et al., 2021).

### 2.4. Whole cell lysate preparation and Immunoblotting

The whole cell lysates preparation and western botting was performed as described by Ghosh et al., 2021 (Ghosh et al., 2021).

### 2.5. shRNA transduction

Second generation of lentiviral transfection system was used to generate viral particle containing shRNA using PEI reagent (Polysciences) as described by Paul et al., 2018 (Paul et al., 2018).

### 2.6. Tumor xenograft model

Two sets each of 20 BALB/c mice (4-6 weeks, 20-25 gm) after acclimatization were injected 2×10^6^ cells CT26-or 4T1 cells/mice. After observation of palpable tumors the animals in each group were randomly segregated into four sub-groups (N=5) for treatment with vehicle or PFBL (100 mg/kg) or 5FU (10 mg/kg) or 5FU+PFBL for CT26 tumors for two weeks. The 4T1 animal groups were treated with the vehicle, PFBL (50 mg/kg), DOX (5 mg/kg) or DOX+PFBL for every alternate day for two weeks. Mice were euthanized, tumors were excised and weighed. The animal experiments were performed per permission from Institutional Animal Ethics Committee (IACE), Bose Institute, Kolkata (Ref. No. IAEC/BI/002/2021).

### 2.7. In vivo metastasis model

For each lung metastasis model, 30 BALB/c mice were injected through tail vain with 2×10^5^ cells (CT26-or 4T1) cells/mice and randomly segregated into three groups and treated with either the vehicle, 25 mg/kg PFBL and 50 mg/kg PFBL every alternate day for 4 weeks. At the end of the experiment, the mice were euthanized, and the lungs were collected for further analysis. All the animal experiments were performed with prior permission from IACE, Bose Institute, Kolkata (Ref. No. IAEC/BI/002/2021).

### 2.8. H & E staining

H & E staining was performed as described by Ghosh et al., 2021 (Ghosh et al., 2021).

### 2.7. Statistical analysis

All the graphs presented were plotted using GraphPad prism 5.03 software. One-way ANOVA and Tuckey’s HSD pot-hoc analysis was performed to find out statistically significant differences in group means. Values with a significance level below 0.05 (P<0.05) were deemed statistically significant. All the statistical analysis were done using R version 4.2.3.

## 3. Results and Discussion

### 3.1. PFBL regresses CT26 and 4T1 cell induced solid tumor in BALB/c mice

Mice-bearing solid tumors were treated with PFBL alone or in combination with 5FU, or DOX as described in the Materials and Methods. After completion of the treatment tumors were excised and measured. Unlike the control/untreated groups which grown over time, there was a gradual decrease in tumor mass in both CT26 and 4T1 groups treated with PFBL alone or in combination with DOX or 5FU [Fig. 1A-C]. The H & E staining of tissue sections of these tumors indicated the clearer regions in treated samples compared to denser regions in vehicle-treated controls [Fig. 1D(i)-(ii)]. Unlike the CT26 group, elevated levels of cleaved PARP1 and caspase 3 in the immunoblots of the tumor lysates indicated the predominant role of apoptosis in 4T1-tumors upon treatments [Fig. 1E, F(i-ii)]. PFBL or its 5FU combination induced ∼3-fold increase of PARP1 cleavage for CT26 tumor. The increase in the cleaved caspase level was ∼2-fold and 1.5-fold for PFBL and 5FU combination, respectively [Fig. 1 E, F (i-ii)]. For 4T1 tumor, PARP1 cleavage was increased by ∼6-fold and ∼8-fold by PFBL and its combination treatments with DOX. In the same treatments the cleaved caspase level was increased by ∼2-fold and 3.5-fold, respectively [Fig.1 E, F (i-ii)]. In stark contrast, PFBL treatments induced autophagy in CT26 tumor as indicated by ∼10-fold increase in LC3II levels although the combination treatments with 5FU was less effective (∼6-fold) [Fig.1 E, F (iii)]. There was about ∼2-fold increase in LC3II level in 4T1 samples treated with PFBL alone or in combination with DOX [Fig. 1E, F(iii)]. Therefore, PFBL regressed CT26 and 4T1 tumors majorly by induction of autophagy and apoptosis, respectively. Furthermore, treatments with PFBL alone, and in combination with 5FU or -DOX increased E-cadherin levels (> 2-folds) while decreasing the N-cadherin levels in both types of tumors by > 2-folds [Fig. 1E, F(iv-v)]. PFBL also reversed the EMT (Brabletz et al., 2018) in both types of the tumors [Fig.1E, F(iv-v)]. Notably, the PFBL dosages used were well tolerated in the healthy mice per monitoring their weights, blood serum enzyme (ALP, SGOT, SGPT), and spleen size assessment (Supplementary Fig. S1).

**Figure 1.**
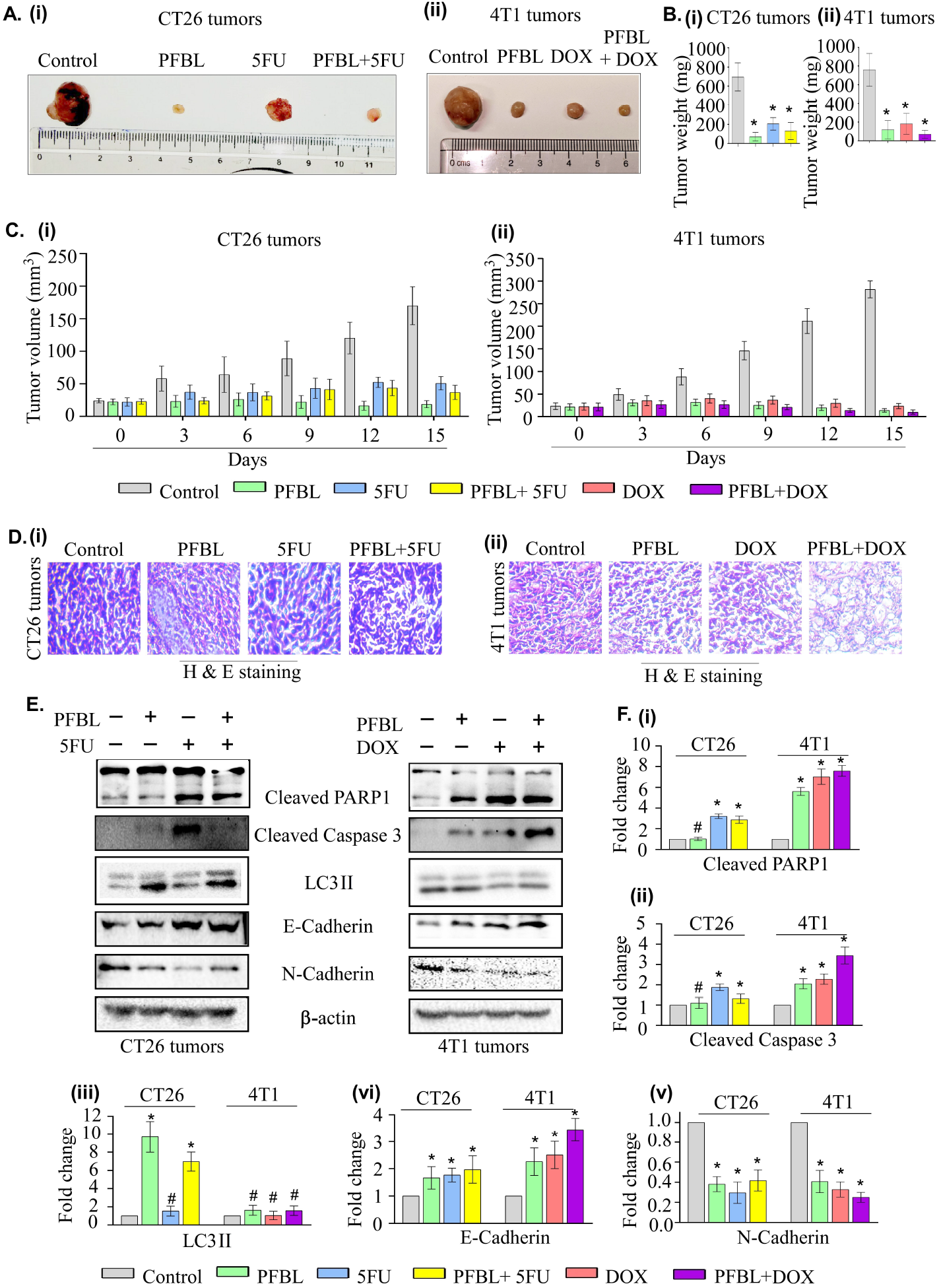
PFBL suppressed CT26 and 4T1 cell induced solid tumor in BALB/c mice models. [A] Representative image of tumors excised from BALB/c mice after treatment as indicated. [B] Bar graph representing the weights of the tumors isolated from the mice. [C] Representative bar graph showing the changes in tumor volumes during the treatment period. [D] Representative images of haematoxylin and eosin stained tumor sections. [E] Immunoblots of whole cell lysate extracted from tumor samples representing different apoptotic, autophagic, and EMT markers. [F] Densitometric quantitation of bands shown by bar graphs normalised using β-actin level. All in comparison with the control group, “#” and “*” represents P value > 0.05 and P value <0.05 respectively.

### 3.2. PFBL suppressed lung metastasis of CT26 and 4T1 cells in BALB/c mice

In vivo anti-tumor activity and changes in EMT marker levels suggested the possibility of an anti-metastatic potential of PFBL [Fig. 1].The metastasis models were developed though injection of CT26 and 4T1 cells through the tail vein as described in the materials and methods. We tested the effect of two different dosages of PFBL in these models. After four weeks of treatment, the mice were euthanized, and the lungs were isolated. For both the lung metastasis model, both in size and weights of lungs, treated with 25 mg and 50 mg of PFBL per kg of mice were smaller than the corresponding vehicle treated group [Fig. 2A-B]. PFBL reduced the size/mass of the lungs by ∼2-fold and the number of lung nodules by >2-fold with increasing dosages of PFBL [Figs. 2A-B]. In the CT26-induced model, the groups treated with the vehicle of PFBL (25 mg/kg and 50 mg/kg body weight of mice) exhibited 76, 37, and 20 nodules on average, respectively, whereas in the 4T1 induced model group the vehicle, 25 mg/kg and 50 mg/kg treated group, on average exhibited 67, 18, and 11 nodules, respectively [Fig. 2C(i)-(ii)]. The H & E staining of lung tissue sections showed larger dense regions in mock-treated groups; the size of the dense regions decreased in samples treated with increasing PFBL concentrations [Fig. 2D]. PFBL showed similar results on EMT markers in HCT116 and MCF7 cells [Supplementary Fig. S8 D-G]. Therefore, PFBL efficiently suppressed lung metastasis of CT26 and 4T1 cells in vivo.

**Figure 2.**
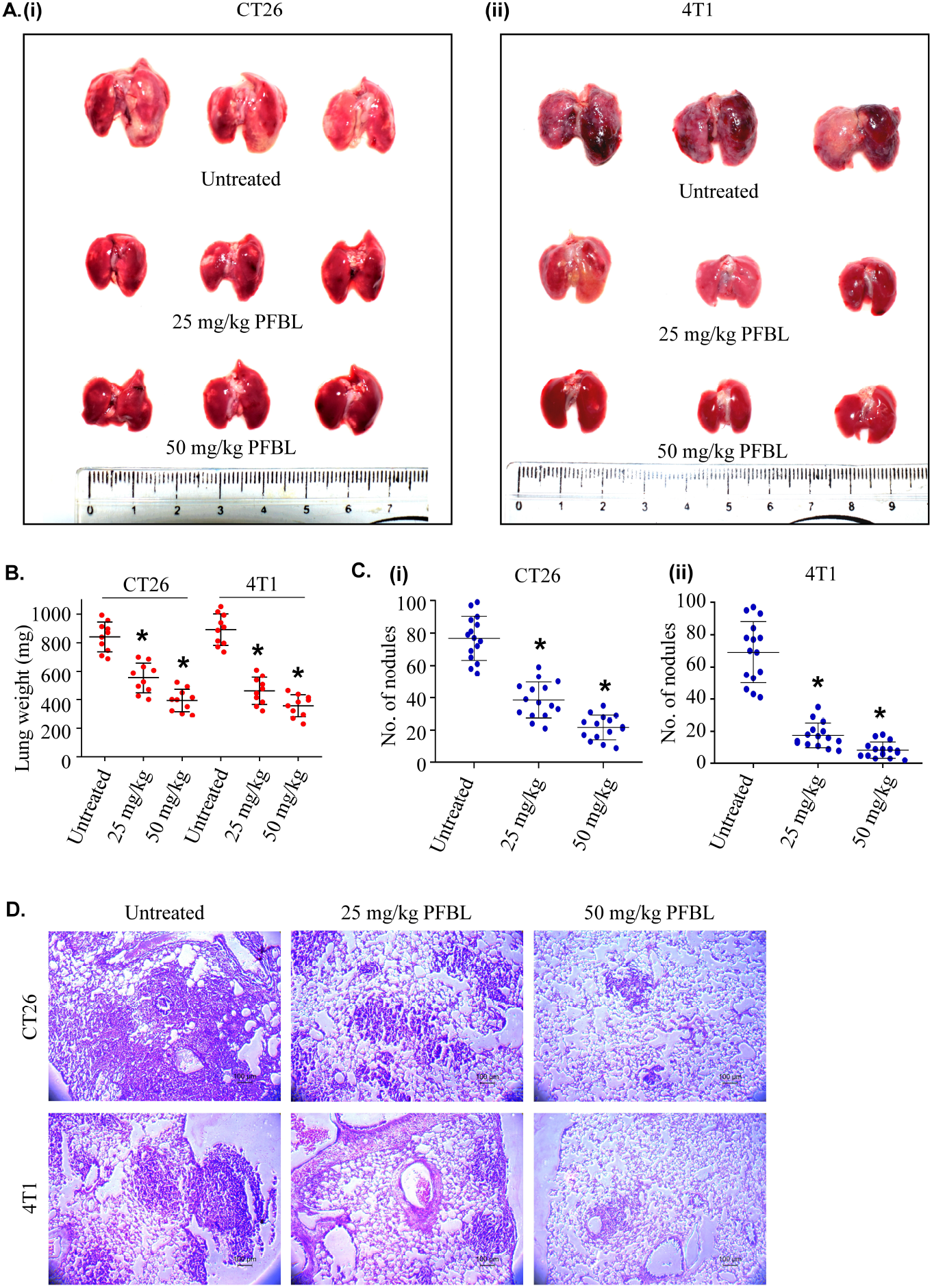
PFBL suppressed lung metastasis of CT26 and 4T1 cells in BALB/c mice model. [A] Representative images of lungs nodules after treatment indicated. [B] Scatter plot representing lung weight of each experimental group. [C] Scatter plot representing the number of lung nodules in each experimental group after as indicated. [D] Representative images of H&E stained lung sections. All in comparison with the untreated control group, “#” and “*” represents P value > 0.05 and P value <0.05 respectively.

### PFBL induced distinct death mechanism in MCF7 and HCT116 cells

Sensitivity of MCF7, HCT116 and NKE cells were tested and the IC50 values were determined (Supplementary Fig. S2). PFBL dose-dependently induced MCF7 and HCT116 cells to arrest in the G0/G1 phase estimated by FACS after propidium iodide (PI) staining [Fig. 3 A(i)-(ii)] (Supplementary Fig. S3). Consistently immunoblots of whole cell lysates (WCLs) of MCF7 and HCT116 cells pre-treated with 40 μg/ml and 50 μg/ml PFBL corresponding their IC50 values for increasing durations indicated significant decrease in the CDK4 and Cyclin D1 levels in both cell types [Fig. 3C-D(i)-(ii)].

**Figure 3.**
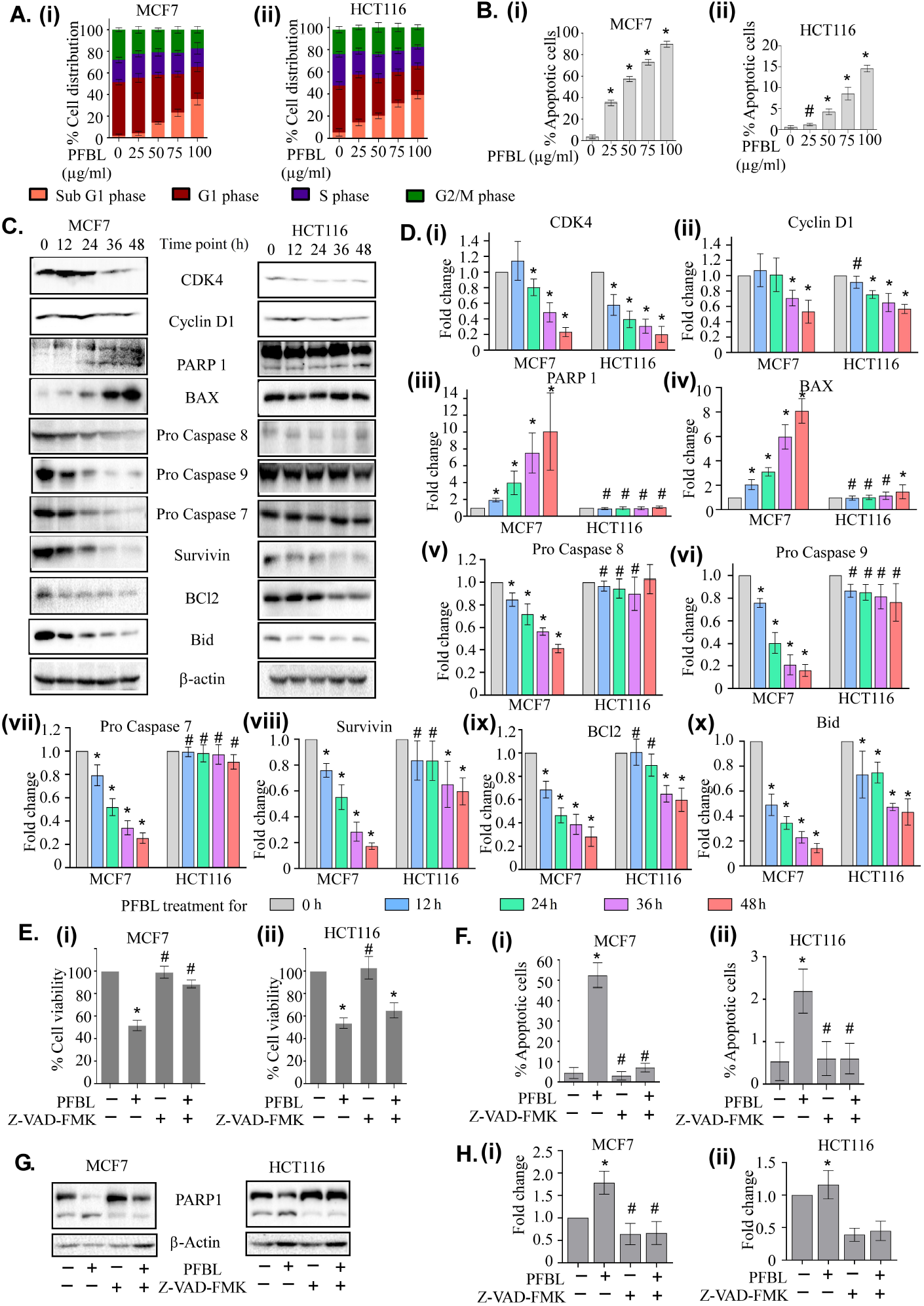
PFBL arrests cells at G0/G1 phase and induces apoptosis. [A] Bar diagram representing the distribution of cells in different cell cycle phases upon treatments as shown. [B] The quantitation of FACS assay representing the status of FITC-Annexin-V and/or PI positive cell population upon treatment as shown in a bar graph. [C] Immunoblots of lysates off cells pre-treated with PFBL for different duration with different cellular apoptotic markers. β-actin level was determined as an internal loading control. Densitometric quantitation of bands relative to β-actin is shown by bar graphs [D] (i)-(x). [E] Bar graph representing the viability (MTT assay) of cells pre-treated with or without pan-caspase inhibitor Z-VAD-FMK for 3 h followed by PFBL treatment. [F] Bar graph representing the percentage of apoptotic cell population cells pre-treated with or without pan-caspase inhibitor Z-VAD-FMK for 3 h followed by PFBL treatment. [G][H] immune blots and corresponding bar graphs representing the fold change of PARP1 level. β-actin level was taken as an internal loading control and measured by densitometric quantitation after Z-VAD-FMK and PFBL treatment. All data represents three independent experimental observations. Significant difference from control group; “#” and “*” represents P value > 0.05 and P value <0.05 respectively.

PFBL dose-dependently induced MCF7 and HCT116 cell death as estimated by FACS after FITC-ANNEXIN-V and propidium iodide (PI) staining [Fig. 3 B(i)-(ii)] (Supplementary Fig. S4). Immunoblots with these WCLs were performed to better understand the molecular pathway of deaths in these cells. In contrast to HCT116 cells, a significant change in the levels of markers of apoptotic signaling pathways, such as cleavage in the PARP1, pro-caspase 7, pro-caspase 8, and pro-caspase 9 molecules including Bcl2, Bid, Survivin, and Bax were apparent in MCF7 cells in consistent with in vivo results [Fig. 3 C-D, Fig. 1]. To verify the significance of apoptosis, these cells pre-treated with pan-caspase inhibitor Z-VAD-FMK were exposed to PFBL for analysis by FACS [Supplementary Fig. S5], MTT assay, and immunoblotting. Pre-treatment with Z-VAD-FMK provided only ∼3% protection to HCT116 cells at 40 μg/ml PFBL (IC50=38.8 μg/ml) [Fig. 5E (ii)-H (ii)]. PFBL at 50 μg/ml (IC50=52.23 μg/ml) killed ∼10% of MCF7 cells when pre-treated with Z-VAD-FMK [Fig. 3 E (i)] down from ∼90% death [Fig. 3 E-H]. In agreement with in vivo results, this data indicated apoptosis as the key mechanism of death in MCF7 cells. Alteration of mitochondrial transmembrane potential (MTP) detected by shifting of JC-1 staining from red to green indicated the cause of ROS production in this process (Supple Fig. S6) (Sivandzade et al., 2019).

Apoptosis apparently played relatively a minor role in the death of HCT116 cells [Fig. 3 E(i)-H(i), Fig. 1]. Guided by the in vivo results we investigated the involvement of autophagy in this case [Fig. 1]. We tested the cleavage of LC3II proteins as an autophagy marker. In agreement with this and in vivo results of immunoblots, PFBL-pre-treated cells showed ∼8-fold and ∼2-fold increase in LC3II level in HCT116 and MCF7 cells, respectively [Figs. 4A, B(i), Fig. 1, Supplementary Fig. S7]. To check the involvement of the mammalian target of rapamycin (mTOR) pathway in PFBL-induced autophagy, different upstream and downstream players of this pathway were tested (Zou et al., 2020). The downregulation of phosphorylation level of AKT, 4EBP1, and S6K with the upregulation of phosphorylation level of AMPK (at T172) indicated mTOR inhibition [Fig. 4 B(ii)-(iv)]. PFBL induced relatively more AMPK phosphorylation in HCT116 cells than MCF7 cells [Fig. 4 B(v)]. To verify the significance of autophagy, cells were pre-treated with autophagy inhibitor chloroquine before treatment with PFBL followed by MTT assay [Fig. 4 C(i)-(ii)]. Chloroquine pre-treatment provided ∼90% protection from toxicity induced by PFBL, indicating the significant role of autophagy in PFBL-induced death of HCT116 cells [Fig. 4 C(i)], where chloroquine pre-treatment provided ∼12% protection [Fig. 4 C(ii)] indicating a minor role of autophagy upon PFBL treatment in MCF7 cells. Differential sensitivity of MCF7 cells to autophagy could be due to their mono-allelic deletion of Beclin 1 gene, a critical player in autophagy (Aita et al., 1999). Unlike normal breast epithelial cells, its expression is diminished in human breast epithelial carcinoma cell lines and tissues. Approximately 40% to 75% of cases of sporadic human breast cancer and ovarian cancers exhibits mono-allelic deletion of Beclin1 gene (Aita et al., 1999).

**Figure 4.**
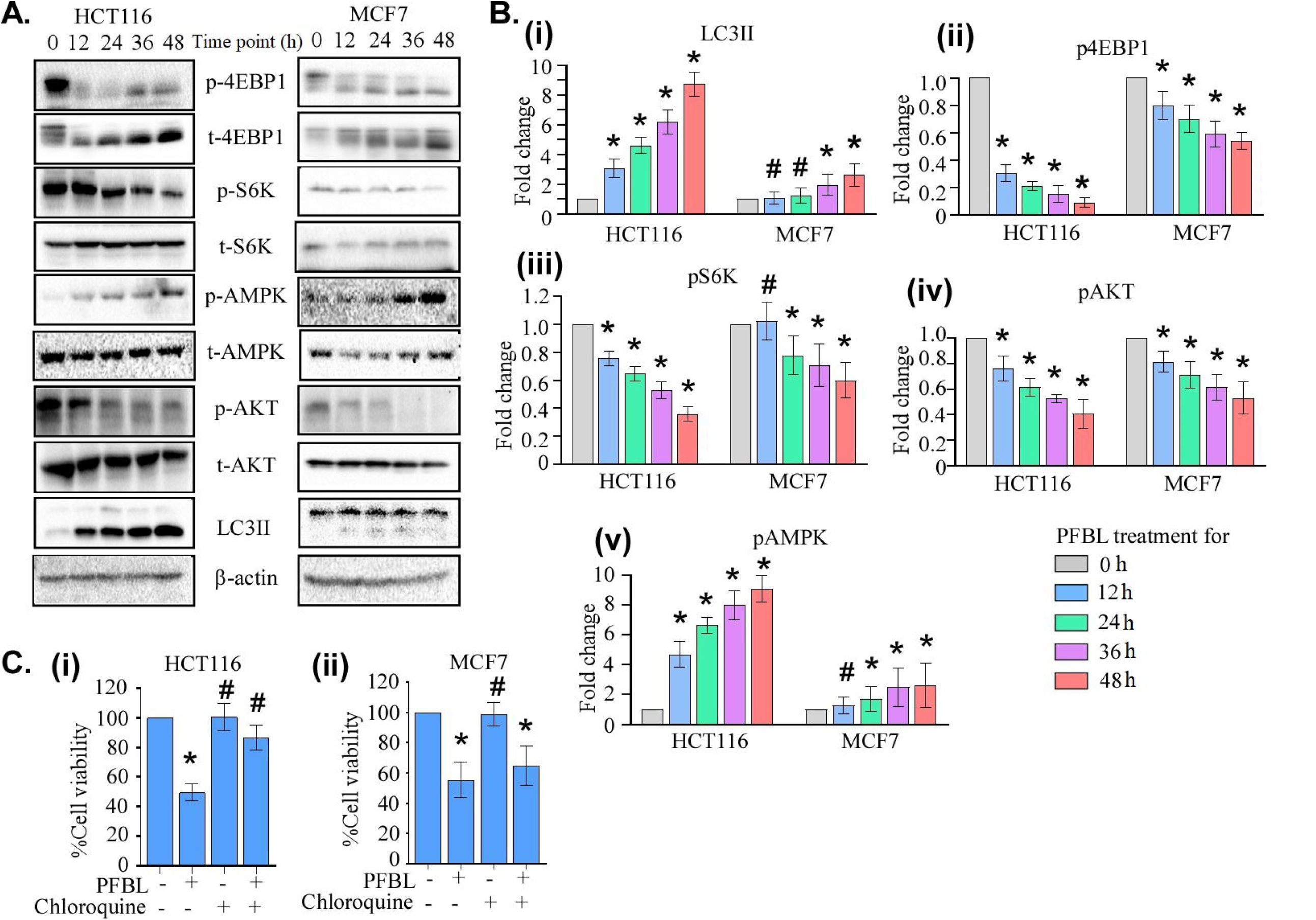
PFBL induces autophagic cell death: [A] Immunoblots of lysates of cells treated with PFBL for different duration. The β-actin level was detected in the same blot as the loading control as indicated. The fold change of signals in the blots was estimated by densitometric quantitation of bands compare to the β-actin level [B]. [C] Cell viability (MTT) assay indicating the role of autophagy (as chloroquine-mediated protection) in PFBL-induced death. All data represents three independent experimental observations. All in comparison with the control group, “#” and “*” represents P value > 0.05 and P value <0.05 respectively.

### 3.4. Role of AMPK in PFBL induced autophagy

AMP-activated protein kinase (AMPK) is a prime sensor and effector of cellular energy metabolism (Hardie, 2007). PFBL induced a dose-dependent increase of AMPK-T172 phosphorylation [Fig. 4 B, C (iv)]. The MTT assay revealed that shRNA-mediated AMPK downregulation increased cells viability along with reduced LC3II levels [Fig 5].

**Figure 5.**
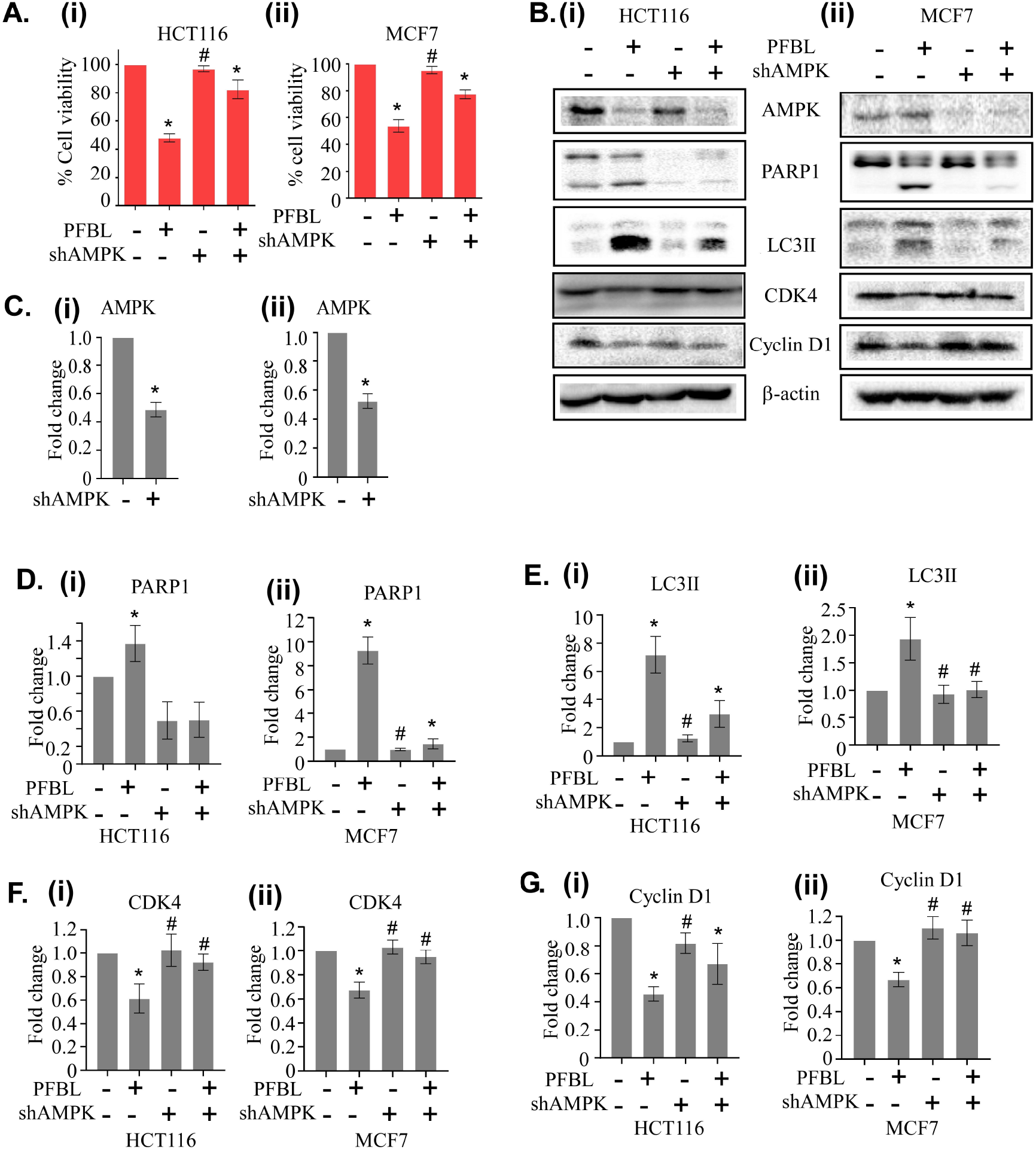
shRNA mediated down regulation of AMPK diminishes the effect of PFBL. [A] Bar graph representing the effect of AMPK downregulation in PFBL-induced cell death determined by MTT assay. [B] Immunoblots representing the effect of AMPK downregulation in PFBL treatment on apoptotic and autophagic markers. The β-actin level was detected as a loading control. The fold change of signals in the blots was estimated by densitometric quantitation of indicated bands compared to the β-actin level as shown in panel [C-G]. All data represents three independent experimental observations. All in comparison with the control group, “#” and “*” represents P value > 0.05 and P value <0.05 respectively.

The involvement of AMPK in cell survival or death, depends on the nature of the stress/signaling and cell type (Villanueva-Paz et al., 2016). AMPK activation induced apoptosis by activation of JNK pathway (Meisse et al., 2002), or Bim mediated apoptosis in breast cancer cells (Concannon et al., 2010) (Jeon et al., 2021). AMPK could block cell cycle progression by stabilizing p53 opposing action by MDMX phosphorylation on p53-Ser342 (He et al., 2014), or by direct phosphorylation of p53-Ser15 (Jones et al., 2005) (Linke et al., 1997). AMPK could induce autophagy during glucose starvation (Herrero[Martín et al., 2009) or through inhibiting mTOR in a TSC2 (Inoki et al., 2003) and Raptor (Gwinn et al., 2008) dependent manner. The mTOR inhibition led to inhibition of cancer cell growth (Zou et al., 2020).

### 3.5. PFBL functions through elevation of intracellular ROS level

ROS can be produced in mitochondria, peroxisome, or endoplasmic reticulum upon exposure to different stimuli including anti-cancer agents (Ghosh et al., 2021). ROS production upon PFBL treatment was examined by DCFDA staining and FACS in cancer cells pre-treated with N-acetylcysteine (NAC) which significantly restored cell viability per MTT assay, along with diminished AMPK phosphorylation comparable to the untreated level [Fig. 6]. Compromised NRF2 antioxidant defence systems make the cancer cells susceptible to enhanced oxidative stress (Perillo et al., 2020). Although a certain level of ROS is required for cancer cell growth, survival, and migration (Covarrubias et al., 2008), higher levels of ROS causes damage to the cellular macromolecules leading to cell death (Halliwell, 2011) (Scherz□Shouval et al., 2007). PFBL was shown to sensitize PCa cells by elevated ROS production through MAO-A activation with simultaneous inhibition of NRF2 antioxidant response involving GSK3β with little effect on normal cells (Ghosh et al., 2021). Normal cells were protected by their fully functional antioxidant defence pathways. ROS also can leads to activation of AMPK (Jeon et al., 2021). Therefore, PFBL-induced elevation of cellular ROS level was the major cause of cell death of cancer cells with defective antioxidant response pathway.

**Figure 6:**
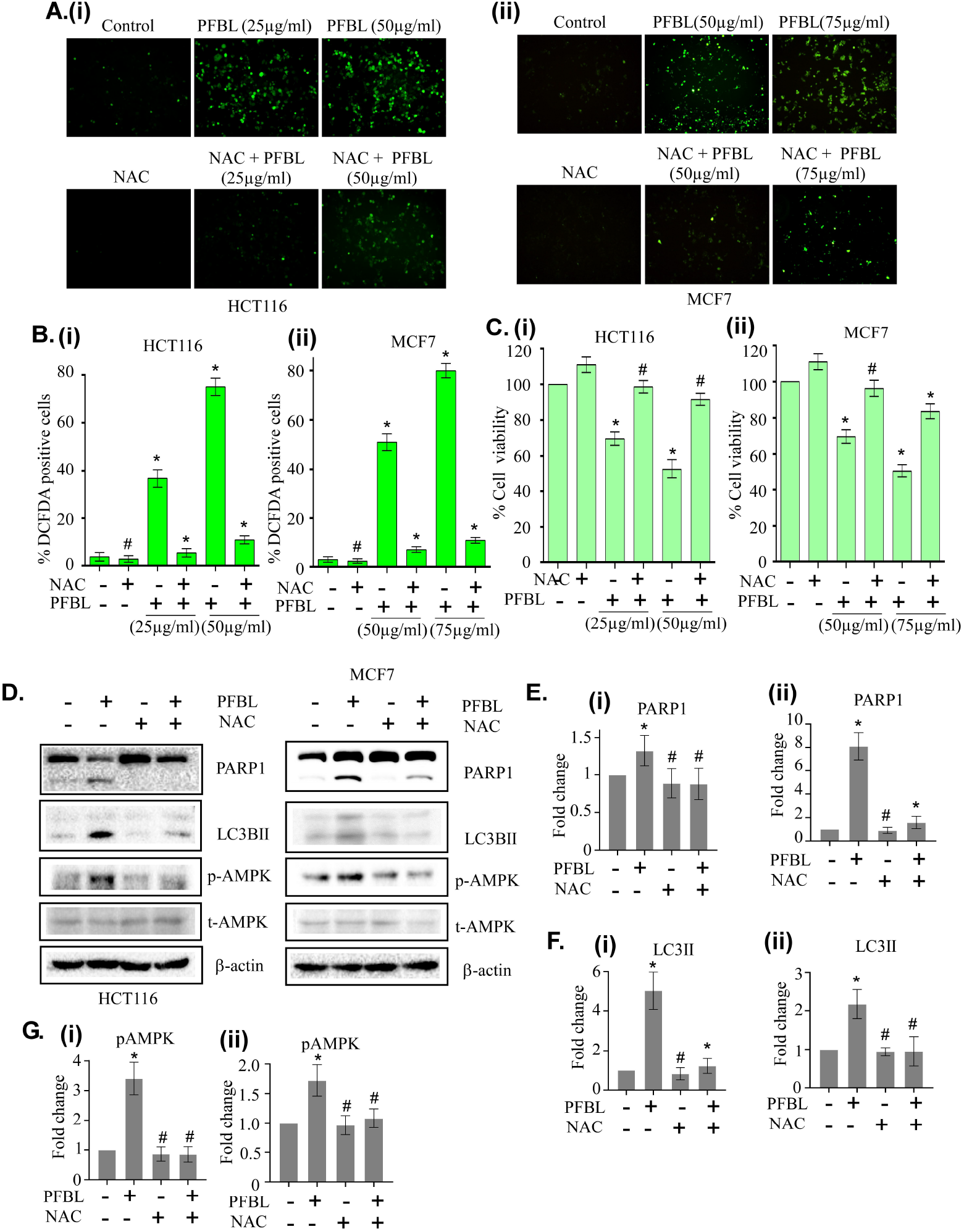
PFBL induced ROS generation is responsible for the death of HCT116 and MCF7 cells. [A] Fluorescence microscopy images of cells treated for 24 h with PFBL with or without NAC pre-treatment for 4 h. The cells were stained with DCFDA before imaging. The DCFDA-positive and viable cell population in the assay are shown in a bar graph in panels [B] and [C], respectively, as indicated. [D] Immunoblots of lysates of PFBL-treated cells with or without pre-treatment with NAC for 4 h with cell markers. The β-actin level was detected as a loading control. The fold change of signals in the blots was estimated by densitometric quantitation of indicated bands compared to the β-actin level as shown in panel [E]. All data represents three independent experimental observations. All in comparison with the control group, “#” and “*” represents P value > 0.05 and P value <0.05 respectively.

This study demonstrates promising therapy potential of PFBL and its capacity to target the cancer cells without affecting healthy cells. PFBL treatment not only regressed CT26 and 4T1-induced solid tumor, but also reduced 5FU- and DOX-induced toxicity when used in combination. Furthermore, PFBL suppressed lung metastasis of CT26 and 4T1 cells in preclinical mice model [Fig. 2]. Detailed investigation revealed HCT116 and MCF7 cell death upon PFBL treatment is predominantly due to autophagic [Fig. 4] and apoptotic cell death [Fig 3], respectively, consistent with in vivo findings [Fig. 1]. PFBL induced ROS production [Fig. 6 A-B] through mitochondrial depolarization (Supplementary Fig. S6) underlies apoptosis in breast cancer cells [Fig. 6] while ROS induced AMPK activation drove in autophagy in the colon cancer cells [Fig. 6].

## Abbreviations

AMPK: AMP-activated protein kinase
BAX: B-cell lymphoma 2 associated x protein
BC: breast cancer
BCL-2: B-cell lymphoma 2
BID: BH3 interacting-domain death agonist
CRC: colorectal cancer
4EBP1: eukaryotic translation initiation factor eIF4E-binding protein 1
GSK3β: Glycogen synthase kinase 3β
LC3B: light chain 3B of microtubule-associated proteins 1A/1B
MAO-A: Monoamine oxidase A
mTOR: mammalian target of rapamycin
NRF2: Nuclear factor erythroid 2-related factor 2
ROS: reactive oxygen species
S6K: ribosomal S6 kinase

## Acknowledgement

This research was supported by Bose Institute, DBT and SERB. Kind supports Profs. Kaushik Biswas and Atin Kumar Mandal are acknowledged. SD was a UGC-adhoc JRF/SRF.

## Author contributions

SD conceptualization, methodology, data curation, all cell-based and animal experiments, immunoblotting, and writing; SG PFBL fraction; SR shRNA cloning and lentiviral transduction; KJ provides animal facility; SCM comments/review; MP conceptualization, supervision, writing, editing.

---

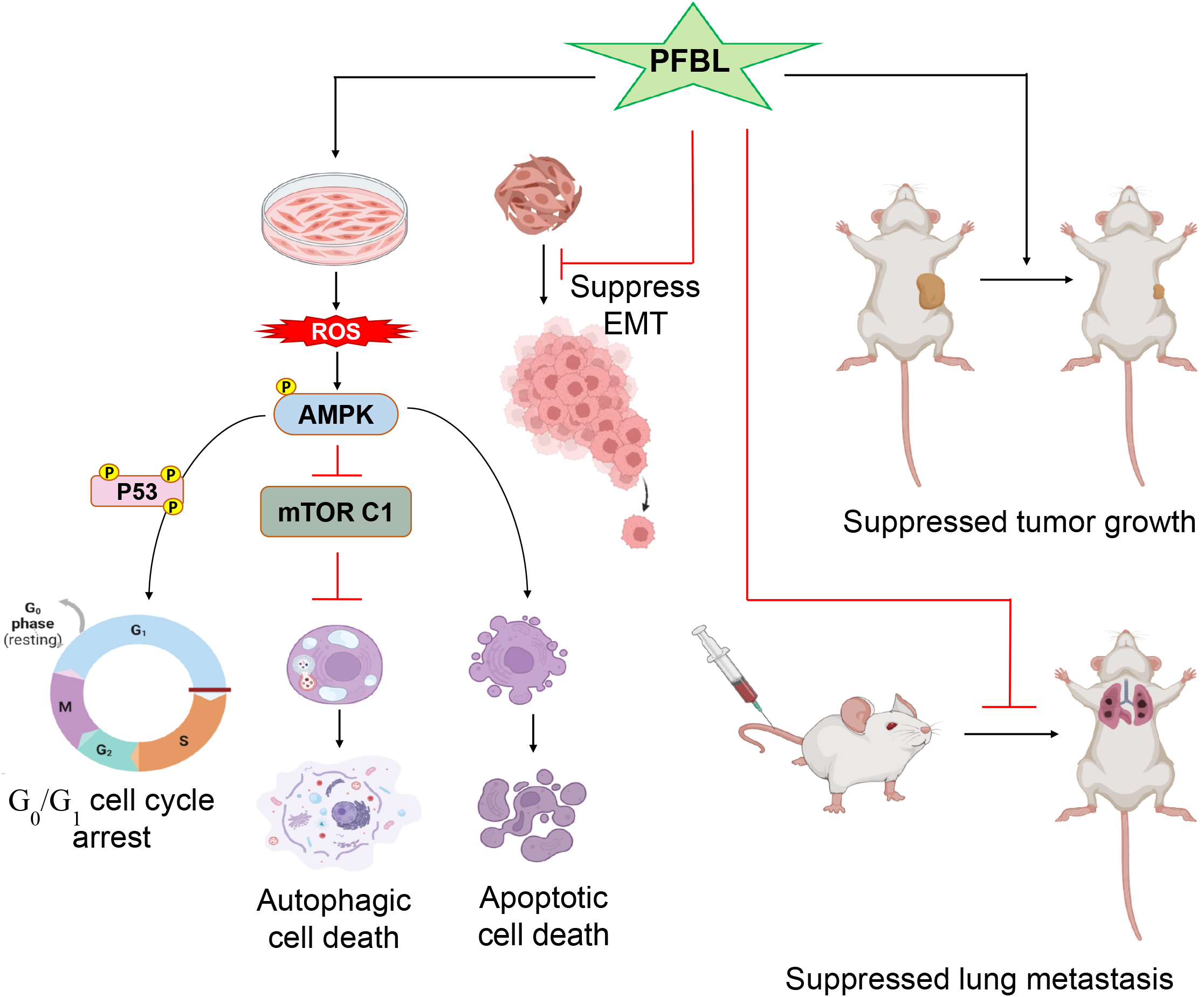

## References

1. Aggarwal, D., Kaushal, R., Kaur, T., Bijarnia, R.K., Puri, S., Singla, S.K., 2014. The most potent antilithiatic agent ameliorating renal dysfunction and oxidative stress from Bergenia ligulata rhizome. Journal of ethnopharmacology 158, 85–93.

2. Agnihotri, V., Sati, P., Jantwal, A., Pandey, A., 2015. Antimicrobial and antioxidant phytochemicals in leaf extracts of Bergenia ligulata: a Himalayan herb of medicinal value. Natural Product Research 29, 1074–1077.

3. Aita, V.M., Liang, X.H., Murty, V., Pincus, D.L., Yu, W., Cayanis, E., Kalachikov, S., Gilliam, T.C., Levine, B., 1999. Cloning and genomic organization of beclin 1, a candidate tumor suppressor gene on chromosome 17q21. Genomics 59, 59–65.

4. Arnold, M., Morgan, E., Rumgay, H., Mafra, A., Singh, D., Laversanne, M., Vignat, J., Gralow, J.R., Cardoso, F., Siesling, S., 2022. Current and future burden of breast cancer: Global statistics for 2020 and 2040. The Breast 66, 15–23.

5. Brabletz, T., Kalluri, R., Nieto, M.A., Weinberg, R.A., 2018. EMT in cancer. Nature Reviews Cancer 18, 128–134.

6. Concannon, C.G., Tuffy, L.P., Weisová, P., Bonner, H.P., Dávila, D., Bonner, C., Devocelle, M.C., Strasser, A., Ward, M.W., Prehn, J.H., 2010. AMP kinase–mediated activation of the BH3-only protein Bim couples energy depletion to stress-induced apoptosis. Journal of cell biology 189, 83–94.

7. Covarrubias, L., Hernández-García, D., Schnabel, D., Salas-Vidal, E., Castro-Obregón, S., 2008. Function of reactive oxygen species during animal development: passive or active? Developmental biology 320, 1–11.

8. De, S., Paul, S., Manna, A., Majumder, C., Pal, K., Casarcia, N., Mondal, A., Banerjee, S., Nelson, V.K., Ghosh, S., 2023. Phenolic phytochemicals for prevention and treatment of colorectal cancer: A critical evaluation of in vivo studies. Cancers 15, 993.

9. Faheem, M.M., Bhagat, M., Sharma, P., Anand, R., 2022. Induction of p53 mediated mitochondrial apoptosis and cell cycle arrest in human breast cancer cells by plant mediated synthesis of silver nanoparticles from Bergenia ligulata (Whole plant). International Journal of Pharmaceutics 619, 121710.

10. Ghosh, S., Dutta, N., Banerjee, P., Gajbhiye, R.L., Sareng, H.R., Kapse, P., Pal, S., Burdelya, L., Mandal, N.C., Ravichandiran, V., 2021. Induction of monoamine oxidase A-mediated oxidative stress and impairment of NRF2-antioxidant defence response by polyphenol-rich fraction of Bergenia ligulata sensitizes prostate cancer cells in vitro and in vivo. Free Radical Biology and Medicine 172, 136–151.

11. Gwinn, D.M., Shackelford, D.B., Egan, D.F., Mihaylova, M.M., Mery, A., Vasquez, D.S., Turk, B.E., Shaw, R.J., 2008. AMPK phosphorylation of raptor mediates a metabolic checkpoint. Molecular cell 30, 214–226.

12. Halliwell, B., 2011. Free radicals and antioxidants–quo vadis? Trends in pharmacological sciences 32, 125–130.

13. Hardie, D.G., 2007. AMP-activated/SNF1 protein kinases: conserved guardians of cellular energy. Nature reviews Molecular cell biology 8, 774–785.

14. He, G., Zhang, Y.-W., Lee, J.-H., Zeng, S.X., Wang, Y.V., Luo, Z., Dong, X.C., Viollet, B., Wahl, G.M., Lu, H., 2014. AMP-activated protein kinase induces p53 by phosphorylating MDMX and inhibiting its activity. Molecular and cellular biology 34, 148–157.

15. Herrero□Martín, G., Høyer□Hansen, M., García□García, C., Fumarola, C., Farkas, T., López□Rivas, A., Jäättelä, M., 2009. TAK1 activates AMPK□dependent cytoprotective autophagy in TRAIL□treated epithelial cells. The EMBO journal 28, 677–685.

16. Inoki, K., Zhu, T., Guan, K.-L., 2003. TSC2 mediates cellular energy response to control cell growth and survival. Cell 115, 577–590.

17. Jeon, H., Huynh, D.T.N., Baek, N., Nguyen, T.L.L., Heo, K.-S., 2021. Ginsenoside-Rg2 affects cell growth via regulating ROS-mediated AMPK activation and cell cycle in MCF-7 cells. Phytomedicine 85, 153549.

18. Jones, R.G., Plas, D.R., Kubek, S., Buzzai, M., Mu, J., Xu, Y., Birnbaum, M.J., Thompson, C.B., 2005. AMP-activated protein kinase induces a p53-dependent metabolic checkpoint. Molecular cell 18, 283–293.

19. Linke, S.P., Clarkin, K.C., Wahl, G.M., 1997. p53 mediates permanent arrest over multiple cell cycles in response to γ-irradiation. Cancer research 57, 1171–1179.

20. Meisse, D., Van de Casteele, M., Beauloye, C., Hainault, I., Kefas, B.A., Rider, M.H., Foufelle, F., Hue, L., 2002. Sustained activation of AMP□activated protein kinase induces c□Jun N□terminal kinase activation and apoptosis in liver cells. FEBS letters 526, 38–42.

21. Paul, S., Ghosh, S., Mandal, S., Sau, S., Pal, M., 2018. NRF2 transcriptionally activates the heat shock factor 1 promoter under oxidative stress and affects survival and migration potential of MCF7 cells. Journal of Biological Chemistry 293, 19303–19316.

22. Perillo, B., Di Donato, M., Pezone, A., Di Zazzo, E., Giovannelli, P., Galasso, G., Castoria, G., Migliaccio, A., 2020. ROS in cancer therapy: the bright side of the moon. Experimental & molecular medicine 52, 192–203.

23. Rajbhandari, M., Wegner, U., Schoepke, T., Lindequist, U., Mentel, R., 2003. Inhibitory effect of Bergenia ligulata on influenza virus A. Die Pharmazie-An International Journal of Pharmaceutical Sciences 58, 268–271.

24. Scherz□Shouval, R., Shvets, E., Fass, E., Shorer, H., Gil, L., Elazar, Z., 2007. Reactive oxygen species are essential for autophagy and specifically regulate the activity of Atg4. The EMBO journal 26, 1749–1760.

25. Singh, A., Tandon, S., Kumar, D., Kaur, T., Kesari, K.K., Tandon, C., 2022. Insights into the cytoprotective potential of Bergenia ligulata against oxalate-induced oxidative stress and epithelial–mesenchymal transition (EMT) via TGFβ1/p38MAPK pathway in human renal epithelial cells. Urolithiasis 50, 259–278.

26. Sivandzade, F., Bhalerao, A., Cucullo, L., 2019. Analysis of the mitochondrial membrane potential using the cationic JC-1 dye as a sensitive fluorescent probe. Bio-protocol 9, e3128–e3128.

27. Sung, H., Ferlay, J., Siegel, R.L., Laversanne, M., Soerjomataram, I., Jemal, A., Bray, F., 2021. Global cancer statistics 2020: GLOBOCAN estimates of incidence and mortality worldwide for 36 cancers in 185 countries. CA: a cancer journal for clinicians 71, 209–249.

28. Villanueva-Paz, M., Cotán, D., Garrido-Maraver, J., Oropesa-Ávila, M., de la Mata, M., Delgado-Pavón, A., de Lavera, I., Alcocer-Gómez, E., Álvarez-Córdoba, M., Sánchez-Alcázar, J.A., 2016. AMPK regulation of cell growth, apoptosis, autophagy, and bioenergetics. AMP-activated protein kinase, 45–71.

29. Zou, Z., Tao, T., Li, H., Zhu, X., 2020. mTOR signaling pathway and mTOR inhibitors in cancer: progress and challenges. Cell & bioscience 10, 31.

