## Supplementary File for "Polyphenol-rich fraction of Bergenia ligulata (Wall.) Engl. sensitizes colon and breast cancer metastasis in vivo: Evidence of cell-type specific mechanism"

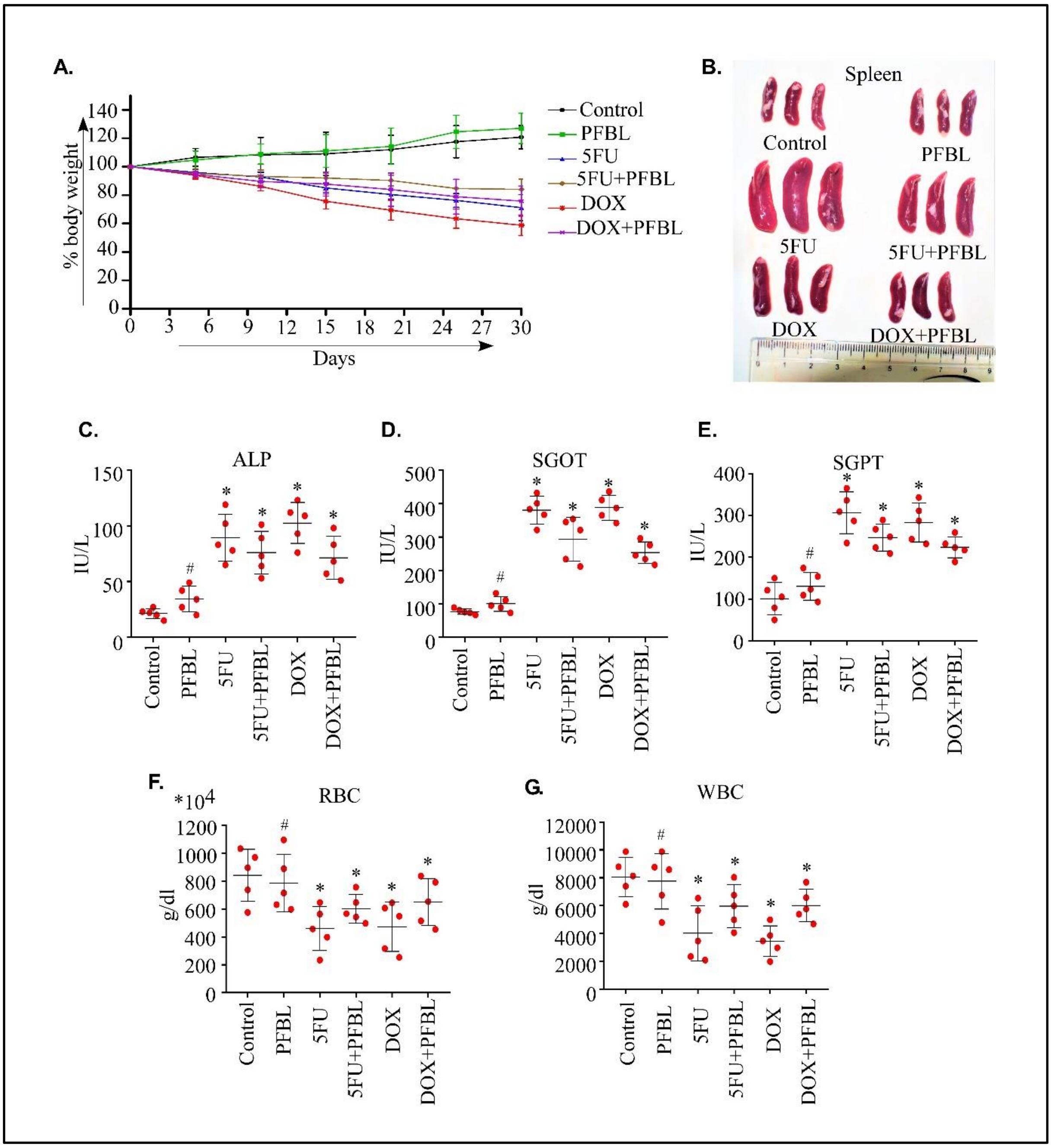


**Supplementary Figure S1:** PFBL is well tolerated in healthy mice and reduces 5FU and DOX-induced toxicity in mice. BALB/c mice were treated with vehicle, or PFBL or 5FU or DOX or their combinations for 30 days. [A] Line graph representing the change in mice body weight during the treatment period. [B] Representative images of the spleens of mice from different experimental groups. The serum alkaline phosphatase (ALP) level [C], serum glutamic-oxaloacetic transaminase (SGOT) level [D], serum glutamic pyruvic transaminase (SGPT) level [E], and the number of RBC [F]and WBC [G] levels were plotted by dot plot. All in comparison with the control group, "#" and "*" represents P value > 0.05 and P value <0.05 respectively.

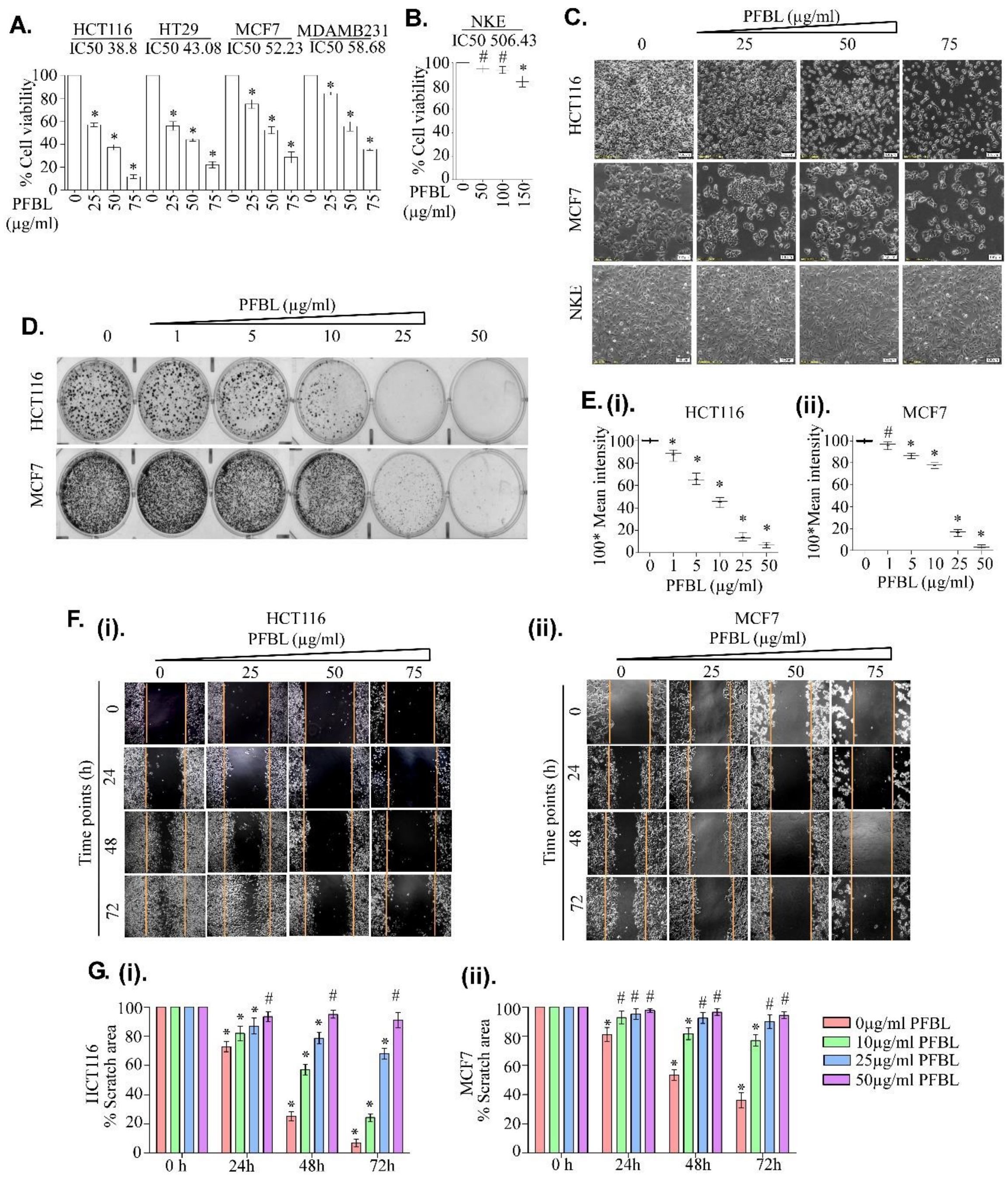


**Supplementary Figure S2:** PFBL sensitized colon and BRC cells but not on NKE cells. [A-B] Bar graph representing the dose-dependent sensitivity of PFBL to human colon cancer (HCT116, HT-29), BRC (MCF7, MDAMB231) and NKE cells determined by MTT assay after 24 h treatment. The IC50 values were shown. [C] Phase contrast images of HCT116, MCF7 and NKE cells treated with increasing concentrations of PFBL for 24 h as indicated. [D-E] Colony-forming ability of cells counted after 24 h treatment. Cells were stained with crystal violet after fixing with formaldehyde. [F] Phase contrast images showing the effect of the treatment on cellular migration potential by wound healing assay. [G] Bar graph representing the estimation of wound healing assay. All data represents three independent experimental observations. All in comparison with the control group, "#" and "*" represents P value > 0.05 and P value <0.05 respectively.

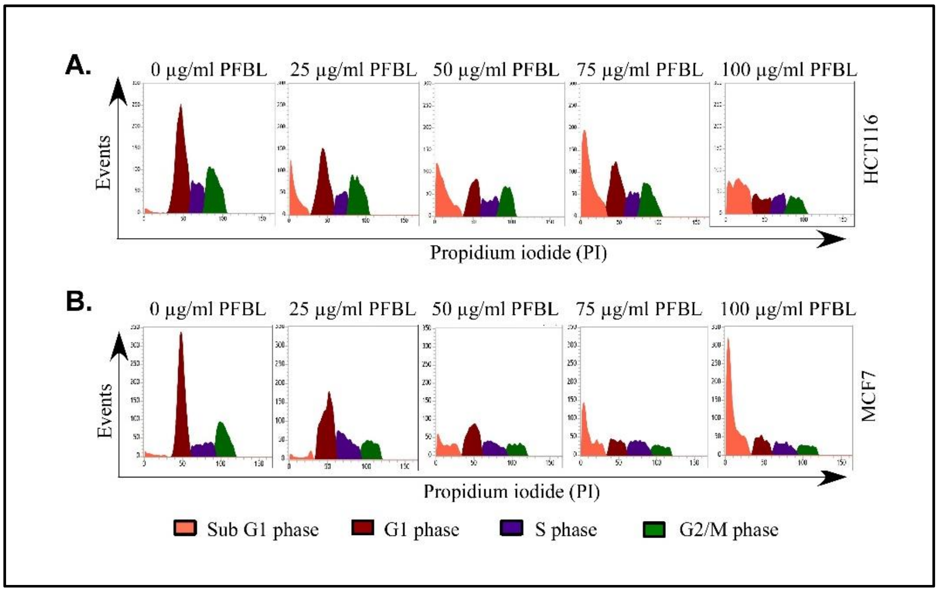


**Supplementary Figure S3:** Histograms showing the effect of different doses of PFBL treatments on cell cycle distribution of HCT116 [A] and MCF7 [B] upon FACS analysis after PI staining.

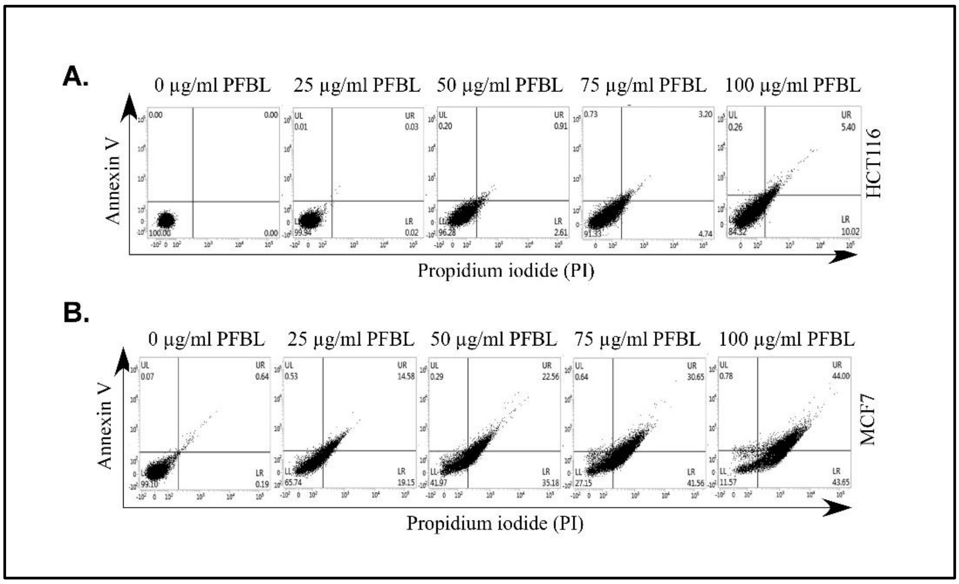


**Supplementary Figure S4:** FACS assay representing the status of FITC-Annexin-V and/or PI positive HCT116 [A] and MCF7 [B] cell population upon treatment with different doses of PFBL.

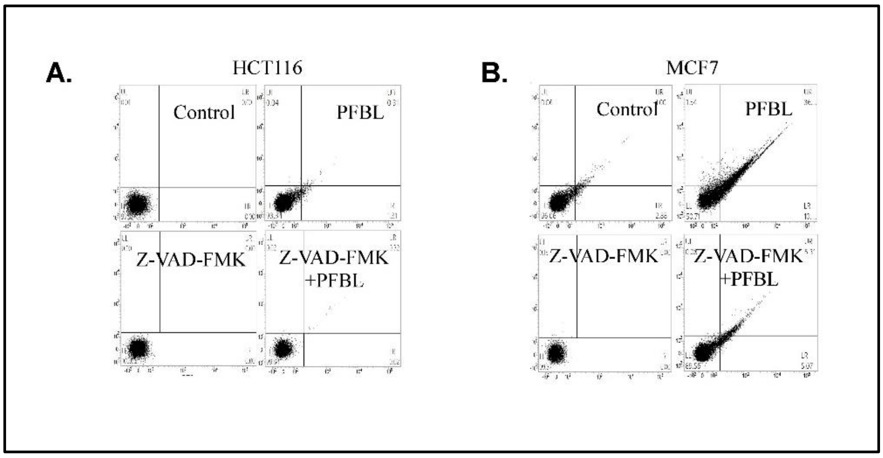


**Supplementary Figure S5:** FITC-Annexin-V and/or PI positive population of HCT116 [A] and MCF7 [B] cells after pre-treatment with or without pan-caspase inhibitor Z-VAD-FMK for 3h followed by 24 h PFBL treatment and assayed by flow cytometry (FACS).

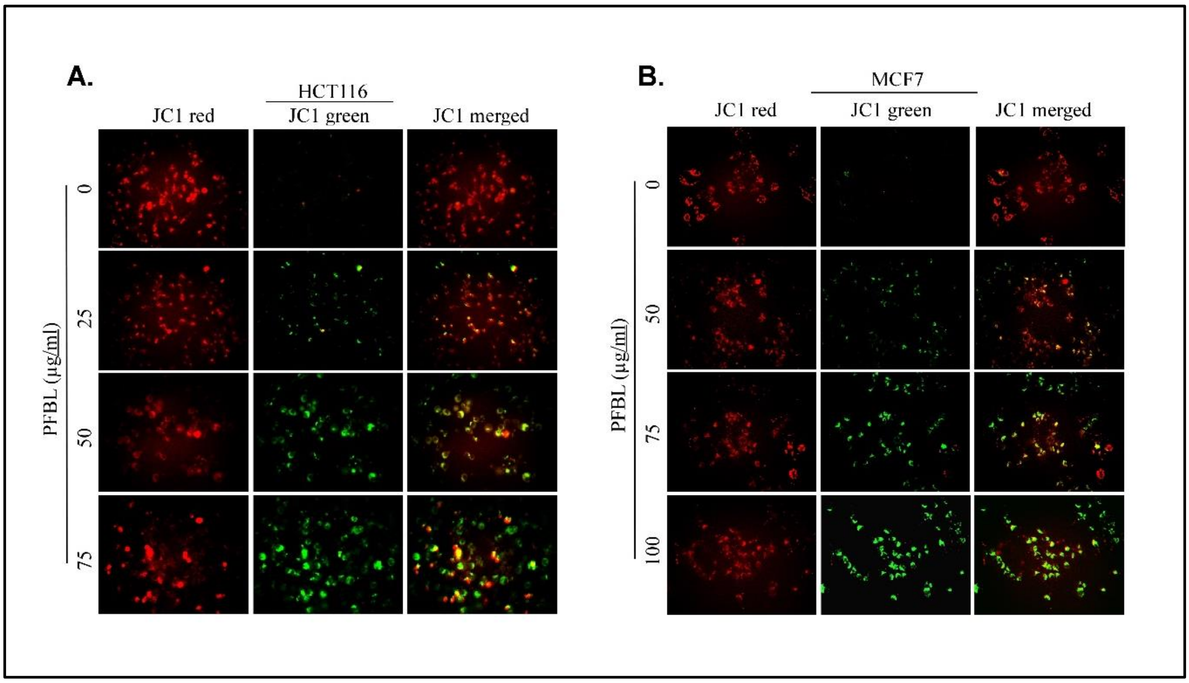


**Supplementary Figure S6:** Fluorescence microscopy images after JC-1 staining representing the change in mitochondrial permeability (from red to green shift) upon treatment with increasing concentrations of PFBL. After treatment, the cells were washed twice with PBS and incubated with JC1 (final concentration of 3 μM) containing serum-free media for 30 minutes at 37°C in a $CO_2$ incubator in the dark. After incubation, cells were washed twice with PBS, and images were captured using a fluorescent microscope. Cells were treated with different concentrations of PFBL followed by JC1 staining. [A] and [B] represents florescence microscopy images of JC1 staining of HCT116 and MCF7 cells respectively.

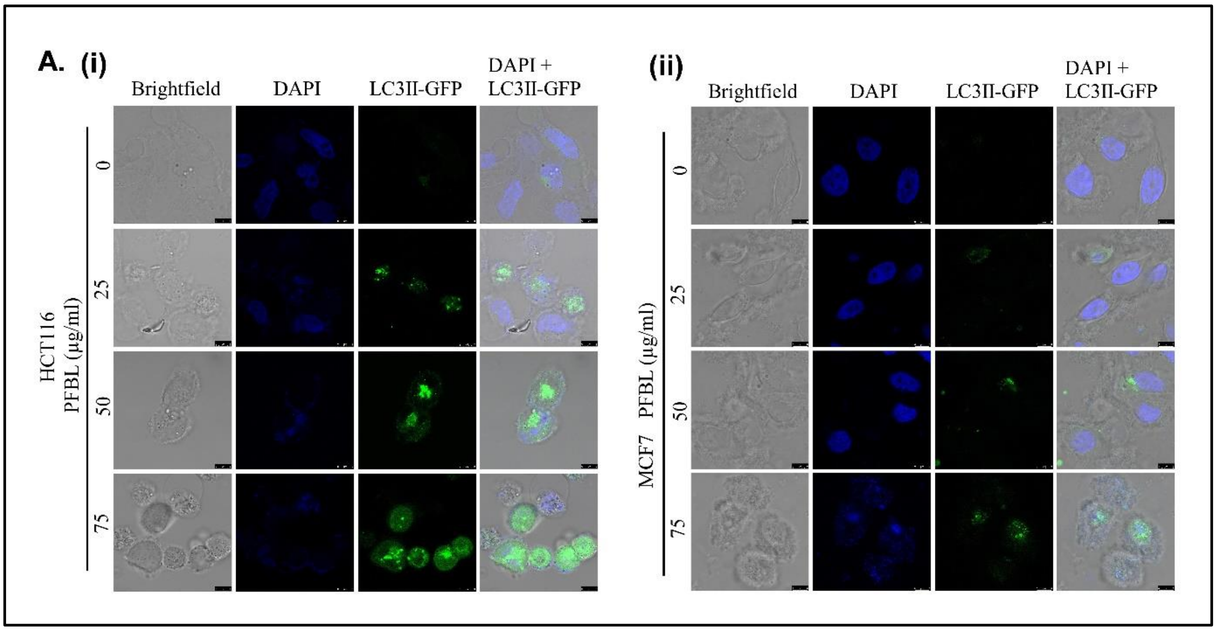


**Supplementary Figure S7:** Confocal microscopy images of LC3II-GFP transfected cells treated with increasing concentrations of PFBL for 24 h followed by DAPI staining.

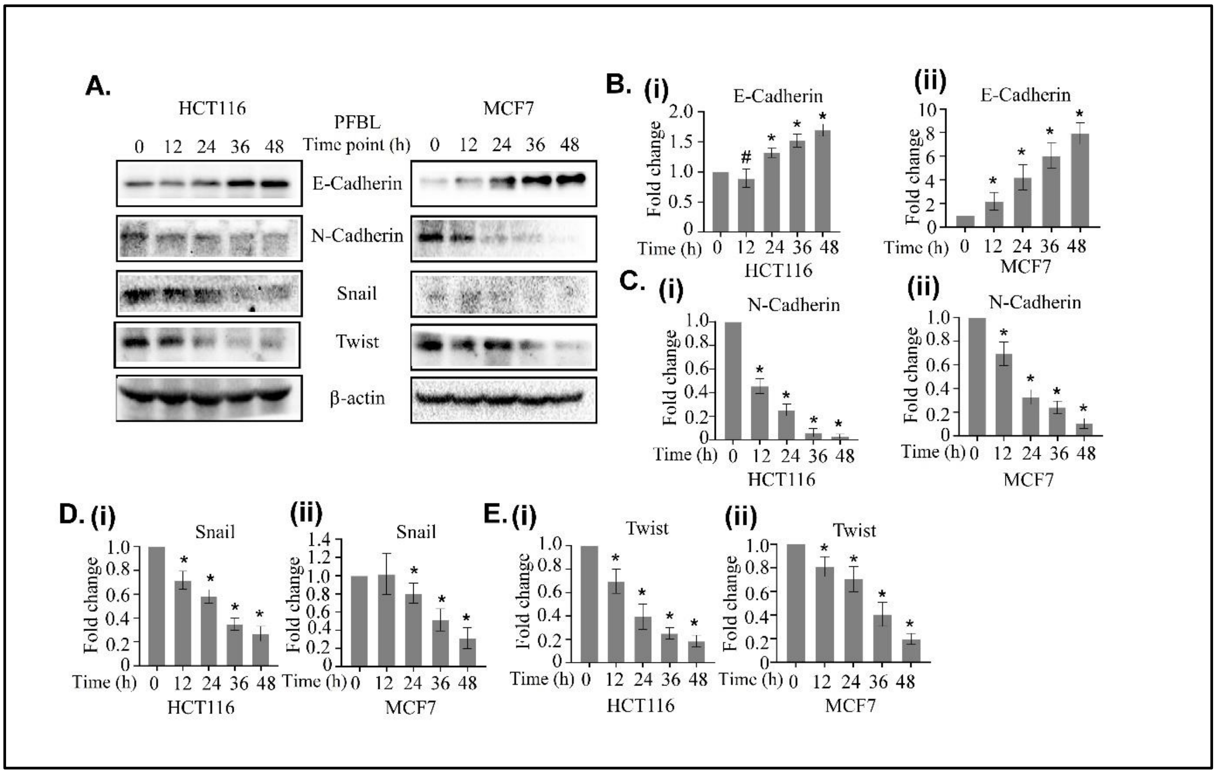


**Supplementary Figure S8:** PFBL inhibits epithelial to mesenchymal transition in HCT116 and MCF7 cells. [A] Immunoblots of lysates of cells treated with PFBL for 24 h for different cellular markers as indicated. The β-actin level was detected as a loading control. [B-E] The fold change of signals in the blots was estimated by densitometric quantitation of indicated bands compared to the β-actin level as indicated. All data represents three independent experimental observations. All in comparison with the control group, "#" and "*" represents P value > 0.05 and P value <0.05 respectively.
